# Evoked pain intensity representation is distributed across brain systems: A multistudy mega-analysis

**DOI:** 10.1101/2020.07.04.182873

**Authors:** Bogdan Petre, Philip Kragel, Lauren Y. Atlas, Stephan Geuter, Marieke Jepma, Leonie Koban, Anjali Krishnan, Marina Lopez-Sola, Mathieu Roy, Choong-Wan Woo, Tor D. Wager

## Abstract

Information is coded in the brain at different scales for different phenomena: locally, distributed across regions and networks, and globally. For pain, the scale of representation is controversial. Although generally believed to be an integrated cognitive and sensory phenomenon implicating diverse brain systems, quantitative characterizations of which regions and networks are sufficient to represent pain are lacking. In this meta-analysis (or mega-analysis) using data from 289 participants across 10 studies, we use model comparison combined with multivariate predictive models to investigate the spatial scale and location of acute pain representation. We compare models based on (a) a single most pain-predictive module, either previously identified elementary regions or a single best large-scale cortical resting-state network module; (b) selected cortical-subcortical systems related to evoked pain in prior literature (‘multi-system models’); and (c) a model spanning the full brain. We estimate the accuracy of pain intensity predictions using cross validation (7 studies) and subsequently validate in three independent holdout studies. All spatial scales convey information about pain intensity, but distributed, multi-system models better characterize pain representations than any individual region or network (e.g. multisystem models explain >20% more of individual subject pain ratings than the best elementary region). Full brain models showed no predictive advantage over multi-system models. These findings quantify the extent that representation of evoked pain experience is distributed across multiple cortical and subcortical systems, show that pain representation is not circumscribed by any elementary region or conical network, and provide a blueprint for identifying the spatial scale of information in other domains.

**Significance Statement:** We define modular, multisystem and global views of brain function, use multivariate fMRI decoding to characterize pain representations at each level, and provide evidence for a multisystem representation of evoked pain. We further show that local views necessarily exclude important components of pain representation, while a global full brain representation is superfluous, even though both are viable frameworks for representing pain. These findings quantitatively juxtapose and reconcile divergent conclusions from evoked pain studies within a generalized neuroscientific framework, and provide a blueprint for investigating representational architecture for diverse brain processes.

**Author Note:** Data storage supported by the University of Colorado Boulder “PetaLibrary”. Research funded by NIMH R01 MH076136, NIDA R01 DA046064 and NIDA R01 DA035484. Lauren Atlas is supported in part by funding from the Intramural Research Program of the National Center for Complementary and Integrative Health, National Institutes of Health (ZIA-AT000030). Marina Lopez-Sola is supported by a Serra Hunter fellow lecturer program. We would like to thank Dr. Christian Buchel for contributing data to this project, and Dr. Marta Čeko for comments and feedback on the manuscript.

## INTRODUCTION

Neuroscience is shaped by the opposition between local and global perspectives. One holds that the brain is modular with distinct functions ascribed to specific brain regions (Fodor 1983) while the other holds that the brain is best understood holistically (Flourens 1824; Lashley 1929). The dialectical opposition of these perspectives is famously underscored by the classic language studies of Broca and Wernicke, which showed unequivocal localization of function, but also identified “disconnection syndromes” resulting from white matter lesions which demonstrate a distributed character for higher order processing (Catani and Ffytche 2005; Mesulam 1990; Wernicke 1874). This dialectic continues to inform contemporary neuroscience in the form of debates over the extent to which the function of a region can be reduced to a single behavior or the converse. This plays out in vision (Kanwisher, McDermott, and Chun 1997; Haxby et al. 2001), emotion (Janak and Tye 2015; Thompson and Neugebauer 2017), and other areas, but is particularly relevant for pain where neurosurgical and neuromodulatory interventions are guided directly by scientific understanding of how pain representations are localized in the brain (Ronald Melzack and Wall 1965; Moisset, de Andrade, and Bouhassira 2016; DeCharms et al. 2005).

Pain is often seen as an intrinsically distributed phenomenon (Baliki and Apkarian 2015; Reddan and Wager 2019). The distributed model is supported by early fMRI data (Becerra et al. 1999; Coghill et al. 1999; Apkarian et al. 2005) and neuroanatomical evidence of widespread nociceptive projections in the brain, including targets in the brainstem and amygdala, (Bernard, Bester, and Besson 1996; Thompson and Neugebauer 2017), SII and insula (Gauriau and Bernard 2004; Craig 2014) and cingulate cortex (Apkarian and Shi 1998; Yamamura et al. 1996). However, this perspective has been criticized for conflating pain-specific representations with a nonspecific salience response (Mouraux et al. 2011; Legrain et al. 2011). Indeed, traditional fMRI analyses cannot distinguish between the information necessary and sufficient for representation from secondary responses cascading across brain systems. Some have argued from a localist perspective that the dorsal posterior insula (Segerdahl et al. 2015b) or anterior midcingulate cortex (Lieberman and Eisenberger 2015) are “fundamental to” or “selective for” pain. This view follows in the tradition of Wilder Penfield, who first provided support for pain localization through electrical stimulation of dorsal posterior insula (Penfield and Jasper 1954; Mazzola et al. 2012) Others suggest that pain may be encoded throughout the brain (Liang et al. 2013) or even be a global brain state (Mansour et al. 2016). Yet, despite two centuries of neuroscientific inquiry into the architecture of brain representations, the best way to characterize pain remains the subject of active debate (Segerdahl et al. 2015b; Davis et al. 2015; Lieberman and Eisenberger 2015; Wager et al. 2016; Segerdahl et al. 2015a; Liberati et al. 2016).

Crucially, most brain studies identify foci of evoked pain response (Apkarian et al. 2005; Becerra et al. 1999; Coghill et al. 1999) or multivariate patterns that predict pain intensity (Wager et al. 2013; Cecchi et al. 2012; Marquand et al. 2010). Neither of these approaches is sufficient to characterize where pain representations are localized, because they do not explicitly test the *spatial scope* of representation--whether any region or pattern is both necessary and sufficient to predict pain experience. Two pioneering studies have investigated the spatial scope of patterns that discriminate painful from nonpainful stimuli (Brodersen et al. 2012; Brown et al. 2011), and provide preliminary evidence in favor of a distributed representation. However, they do not test an exhaustive set of elementary regions or a comprehensive range of spatial scales. In addition, they do not generalize across diverse task conditions or diverse cognitive and nociceptive sources. And finally, they suffer from a number of statistical limitations, including small sample sizes (N=16), which raises questions regarding their reproducibility and generalizability (Button et al. 2013). Nevertheless these studies make important contributions, including advancing multivariate modeling as a method of quantifying unique pain information across brain areas.

Here we investigate the spatial architecture of pain representation using multivariate decoding of pain intensity across a diverse set of physical and psychological pain manipulations and a large multi-study cohort (N=289, 10 studies). We formalize a modular perspective by testing predictive models in each of 486 elementary brain parcels (including cortical areas from (Glasser et al. 2016), and others), and in defined resting state connectivity modules (including 7-network and 32-network scales; (Yeo et al. 2011)). These allow us to test whether any single, isolated region or any single network module (e.g., the ‘salience network’) is sufficient to decode pain intensity. We formalize the distributed, multi-system perspective by testing spatial basis patterns that cross resting-state modular boundaries, including *a priori* multi-system maps and the whole brain. Formal model comparisons at each spatial scale provide a quantitative measure of the extent to which modular features, distributed networks and the whole brain represent pain experience. By repeating this process across a selection of datasets with varying data acquisition and pain manipulations we are able to generalize our findings to novel studies and contexts.

## METHODS

### Participants

This study aggregated data from 10 previously published studies involving 289 participants. Institutional review boards at the University of Colorado Boulder, Columbia University or the ethics committee of the Medical Chamber Hamburg approved these studies. Please refer to Table 1 for details on sample size, age ranges and gender. All subjects are right handed.

**Table 1.** Study participants overview

|  | Sample size | Gender | Mean Age (SD) | Citations |
| --- | --- | --- | --- | --- |
| Study 1 (BMRK4) | 28 | 10F | 25.7 (7.4) | (Krishnan et al. 2016) |
| Study 2 (EXP) | 17 | 9F* | 25.5 | (Atlas et al. 2010) |
| Study 3 (IE2) | 16 | 9F | 25.9 (10.0) | (Jepma et al. 2018) |
| Study 4 (ILCP) | 29 | 16F* | 20.4(3.3)** | (Woo et al. 2017) |
| Study 5 (NSF) | 26 | 9F | 27.8 | (Wager et al. 2013; Atlas et al. 2014) |
| Study 6 (Romantic Pain) | 30 | 30F | 24.5 (6.7) | (López-Solà et al. 2019) |
| Study 7 (SCEBL) | 25 | 11F | 27.4 (8.8) | (Koban et al. 2019) |
| Study V1 (BMRK3) | 33 | 22F | 27.9 (9.0) | (Wager et al. 2013; Woo et al. 2015) |
| Study V2<br>(IE) | 45 | 25F | 24.7 (7.1) | (Roy et al. 2014) |
| Study V3<br>(Placebo) | 40 | 0F | 26 [19,40]*** | (Geuter et al.<br>2013) |

\* gender of one participant unknown, \*\* age of one participant unknown. \*\*\*standard deviation unavailable, range provided instead

### Data

All data in this study have been previously published. Refer to Table 1 for references to publications containing relevant experimental design and data acquisition details and Table 2 for overviews of experimental designs. Data are available as ‘single trial’ statistical contrast maps, hosted at figshare.com and best accessed via the CANlab Single Trials Repository on github (https://github.com/canlab/canlab_single_trials).

**Table 2.** Study stimuli overview

|  | Stimulus Intensity (°C) | Stimulus Duration | Stimulus Location | Thermode Size (mm<sup>2</sup>) | Trials per Subject | Other experimental manipulations |
| --- | --- | --- | --- | --- | --- | --- |
| Study 1 (BMRK4) | 44.5-48.5* | 11s | L forearm, L foot | 16 | 81 | Learned heat-predictive visual cues |
| Study 2 (EXP) | 41.1-47.1 | 11s | L forearm | 16 | 64 | Learned heat-predictive auditory cues |
| Study 3 (IE2) | 48, 49 | 1s | L forearm | 27 | 70 | Learned heat-predictive visual cues |
| Study 4 (ILCP) | 45, 47 | 10s | L forearm | 16 | 64 | Perceived control, learned heat-predictive visual cues |
| Study 5 (NSF) | 40.8-47.0 | 11s | L forearm | 16 | 48 | Masked emotional faces |
| Study 6 (Romantic Pain) | 47 | 11s | L forearm | 16 | 16 | Romantic partner handholding |
| Study 7 (SCEBL) | 48, 49, 50 | 2s | R calf | 27 | 96 | Unlearned visual expectation cue |
| Study V1 (BMRK3) | 44.3, 45.3, 46.3, 47.3, 48.3, 49.3 | 12.5s | L forearm | 16 | 97 | Cognitive self-regulation of pain (reappraisal) |
| Study V2 (IE) | 47.4* | 4s | L forearm | 16 | 48 | Learned heat-predictive visual cues, placebo |
| Study V3 (Placebo) | 46.4* | 20s | L forearm | 30 | 60 | unlearned visual cues placebo |

\*Variable across participants. Mean across subjects shown

Each study involved a series of trials of painful stimuli of varying intensities (40.8C - 50C) applied to the left arm, anticipatory cues, noxious stimuli lasting 1 to 20 sec, and a pain rating period following a variable (jittered) interval (Figure 1A). Additionally many studies included a cognitive or affective manipulation like romantic partner hand holding or manipulations of perceived control over stimulus intensities (Figure 1B). Table 2 details stimulus durations, sites, intensities, and additional protocol details by study. Table 3 provides effect sizes for experimental pain manipulations across studies.

**Figure 1.**
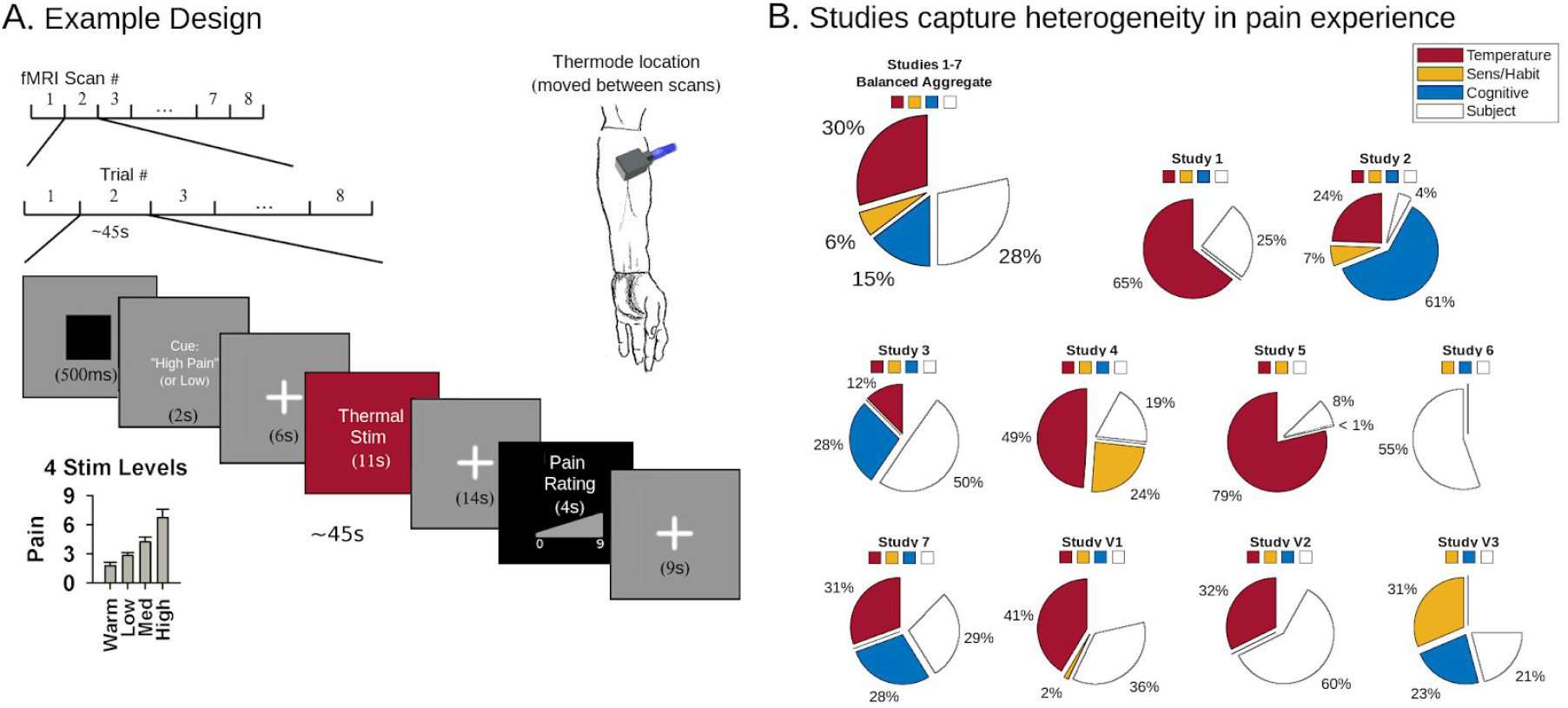
Study design. (A) Each experimental design involves a succession of thermal boxcar stimuli delivered at variable but finite intervals over a succession of fMRI sessions. A cue or task designed to affect pain intensity precedes the stimulus in many designs, for example a pain predictive cue designed to heighten or reduce pain experience. Thermal stimuli vary in intensity and are mostly delivered to the left ventral forearm as illustrated. Cues vary across studies and are uncorrelated with stimulus intensity. After each stimulus subjects rate their pain intensity using a visual scale. For details on each design, refer to Table 2. (B) Pain ratings reflect a variety of influences on pain experience. Wedges show the proportion of variance explained by each factor for each study individually and across all studies (excluding validation studies, balancing subjects across studies). Since fMRI data is quartiled for computational tractability, and z-scored for between study normalization, this data is also quartiled and (in the aggregated data) z-scored prior to regression analyses. The sum of wedges equals model variance explained. Residual model error is not shown, but would merely complete the circles. We do not model subject fixed effects for MVPA but we display them here to better characterize non-experimental sources of pain variability. See Table 3 for standardized regression coefficients. Illustrated arm adapted from Da Vinci’s Vitruvian man (da Vinci c 1492).

**Table 3.**
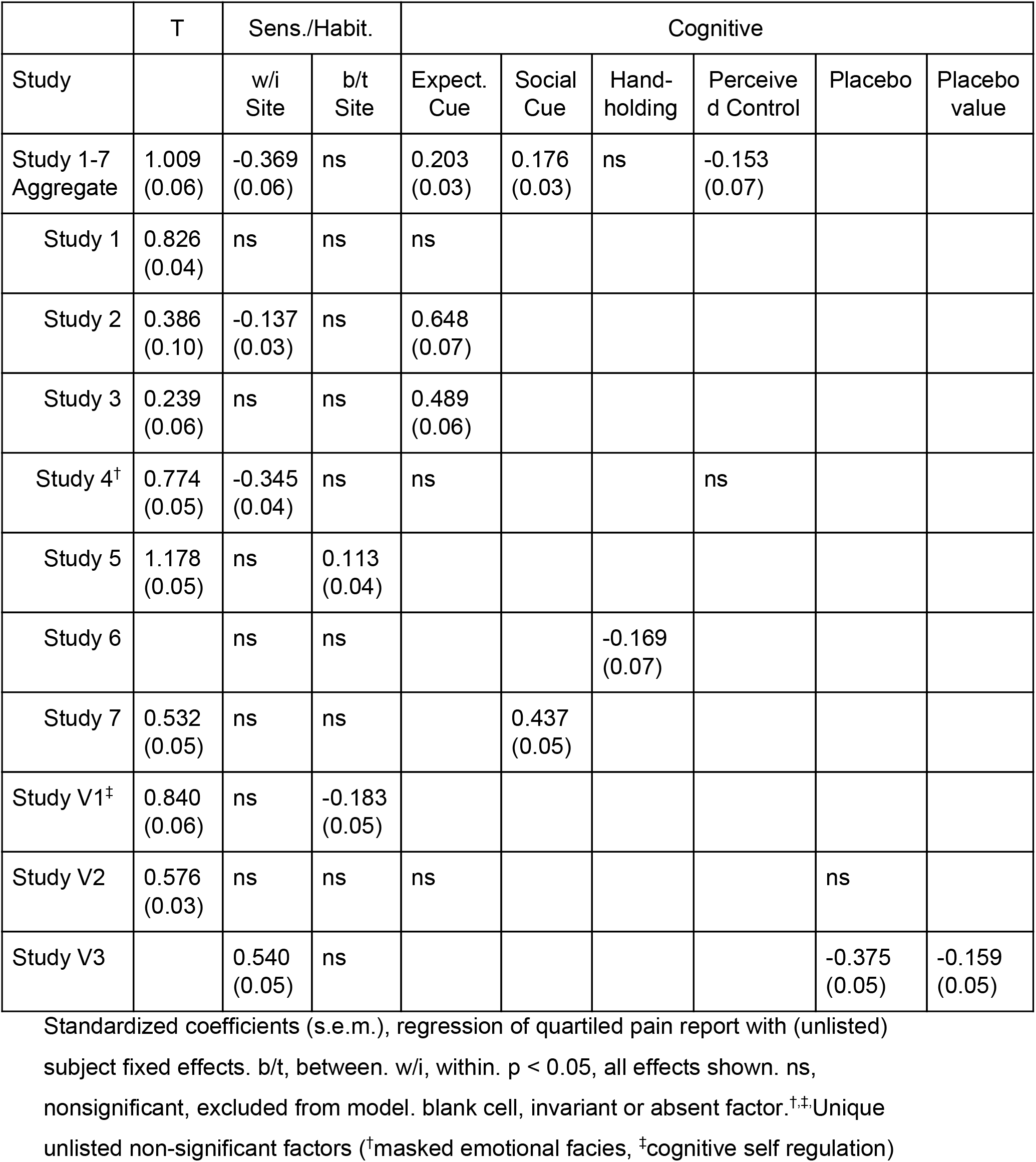
Study behavioral factors overview. Regression model coefficients (Figure 1B) shown.

We perform all of our primary analyses in seven studies (Studies 1-7), and use 3 validation studies (Studies V1-V3) as a post-hoc confirmation of the validity of our inferences. Importantly, the use of validation studies is separate from the use of cross validation, which is used extensively for model fitting and evaluation (see MVPA sections below): validation studies test an estimate while cross validation generates an estimate, and validation studies were left untouched and uninspected until all of our primary analyses had been completed.

### Behavioral data analysis

We model pain report as a function of experimental manipulations and subject fixed effects to provide a descriptive characterization of each study. We consider all experimental manipulations, as well as site specific and non specific sensitization and habituation effects, and then use backwards stepwise regression to select parameters which significantly affect pain report (p < 0.05). All non-significant parameters other than subject effects are removed. For simplicity, this study does not consider nonlinear effects (i.e. interaction effects). All analyses are performed in quartiled data to better inform the subsequent MVPA analyses, which are similarly quartiled.

Partial coefficient of determination (r^2^) quantifies variance explained. Formal definitions in multivariate contexts vary, and the most common are marginal, quantifying the amount of additional variance explained by the inclusion or omission of a variable from a model, e.g. (Anderson-Sprecher 1994), however, when variables have any collinearities they will show some overlap in the outcome variance they can explain which marginal r^2^ calculations omit. We instead adopt the method described by (Scherrer 1984) which splits the difference for collinear independent variables. We take the covariance of the independent variable with predicted pain report (a univariate analysis) and scale it by the standardized regression coefficient of the multivariate model of pain report just described. Computed this way, the individual r^2^ sum to 1, validating their naive interpretation as “percent variance explained”.

### fMRI Acquisition and Preprocessing

Briefly, all data were acquired using 3T MRI scanners, and were motion corrected, had cerebrospinal fluid and white matter signals linearly removed, and were spatially normalized to a standard MNI152 template, except for Study 2 and Study 5, which were acquired at 1.5T and did not have cerebrospinal fluid or white matter signals removed. Please refer to studies referenced in table 1 for further details on data preprocessing.

### General linear model (GLM) analysis

This study uses beta-series analysis (Rissman, Gazzaley, and D’Esposito 2004) to estimate brain response to each evoked noxious stimulus (a single “trial”). The GLM design matrix includes separate regressors for each stimulus and nuisance parameters (e.g. head motion). This study excludes trials with variance inflation factors > 2.5. We thus obtain a separate parameter estimate brain map for stimulus evoked activity for each stimulation (trial). For computational tractability we average these maps within pain intensity quartile after data standardization (see “Data Standardization” below).

### Data Standardization

We remove between-study voxel variability (i.e. the study mean maps), and standardize all single trial images within-trial (mean voxel value is zero, standard deviation of voxel values within trial image is 1). This removes global intensity differences between images, but preserves relative spatial differences within images. Thus, individual images differ from one another in the spatial configuration of their pain response but are matched on net response. Although global intensity information may be task related, it is also affected in idiosyncratic ways by preprocessing decisions, image acquisition parameters, and spurious physiological noise.

We also z-score pain ratings within study to partially control for quantitative differences in the rating scales and calibrations used between studies. This preserves between-subject differences within-study. Following standardization, imaging data and pain ratings are averaged within the pain rating quartiles for each subject. This removes some portion of the overall variance in our data, and all reported r^2^ statistics should be interpreted accordingly as variance explained in quartiled data, but this is necessary for computational tractability. Incidentally, this also partially aligns variability in our brain data with our outcome variable (pain rating), improves the sensitivity of principal component analysis to pain relevant features, and results in better multivariate models when we subsequently employ principal component regression (see MVPA “General Method Description” below).

### Defining hypothesis spaces and model inputs

We define three hypotheses regarding scope of pain representation (modular, *a priori* multisystem areas or full brain) and formalize them using one or more of six hypothesis spaces (explicitly enumerated below) which constrain subsequently trained machine learning algorithms to draw on sources of information from particular brain areas (Figure 2A). The “modular” hypothesis contends that pain intensity information is best captured in a single physiologically coherent brain area, but is agnostic with respect to location and size of the brain area. To formalize this we draw on three brain parcellations which vary in areal size from fine to coarse. (1) Our finest representation, the “regional” parcellation, defines a set of elementary brain areas by combining the Human Connectome Project’s cortical parcellation (Glasser et al. 2016) with individual cerebellar lobules, and thalamic basal ganglia and brainstem nuclear parcellations. (2) A fine resting state network (fRSN) parcellation involving 32 distinct resting state network parcels and (3) a coarse resting state network (cRSN) parcellation using 7 parcels formalize the network hypothesis, including the hypothesis that a salience network response may largely explain the brain’s response to evoked pain. The hypothesis that pain has a “multisystem” representation assumes there is a pain specific configuration of brain activity, localized in a finite but not necessarily contiguous set of brain areas. We formalize this hypothesis as two additional hypothesis spaces. (4) First by identifying *a priori* the subset of the regional parcellation which receives afferent nociceptive information either directly or indirectly, and taking its union to define a hypothesis space (“pain pathways”). (5) Next we take an empirical approach using a reverse inference metaanalysis map for “pain” from neurosynth.org. Finally, the holistic hypothesis assumes that any of a number of sensory or cognitive functions may recruit any one brain area and therefore pain predictive information could be distributed anywhere and everywhere in the brain. (6) Models trained on the entire brain formalize this final hypothesis. Thus, three hypothesis spaces formalize the modular hypothesis, two formalize the multisystem hypothesis and one formalizes the holistic hypothesis. We elaborate on how each space is defined below.

**Figure 2.**
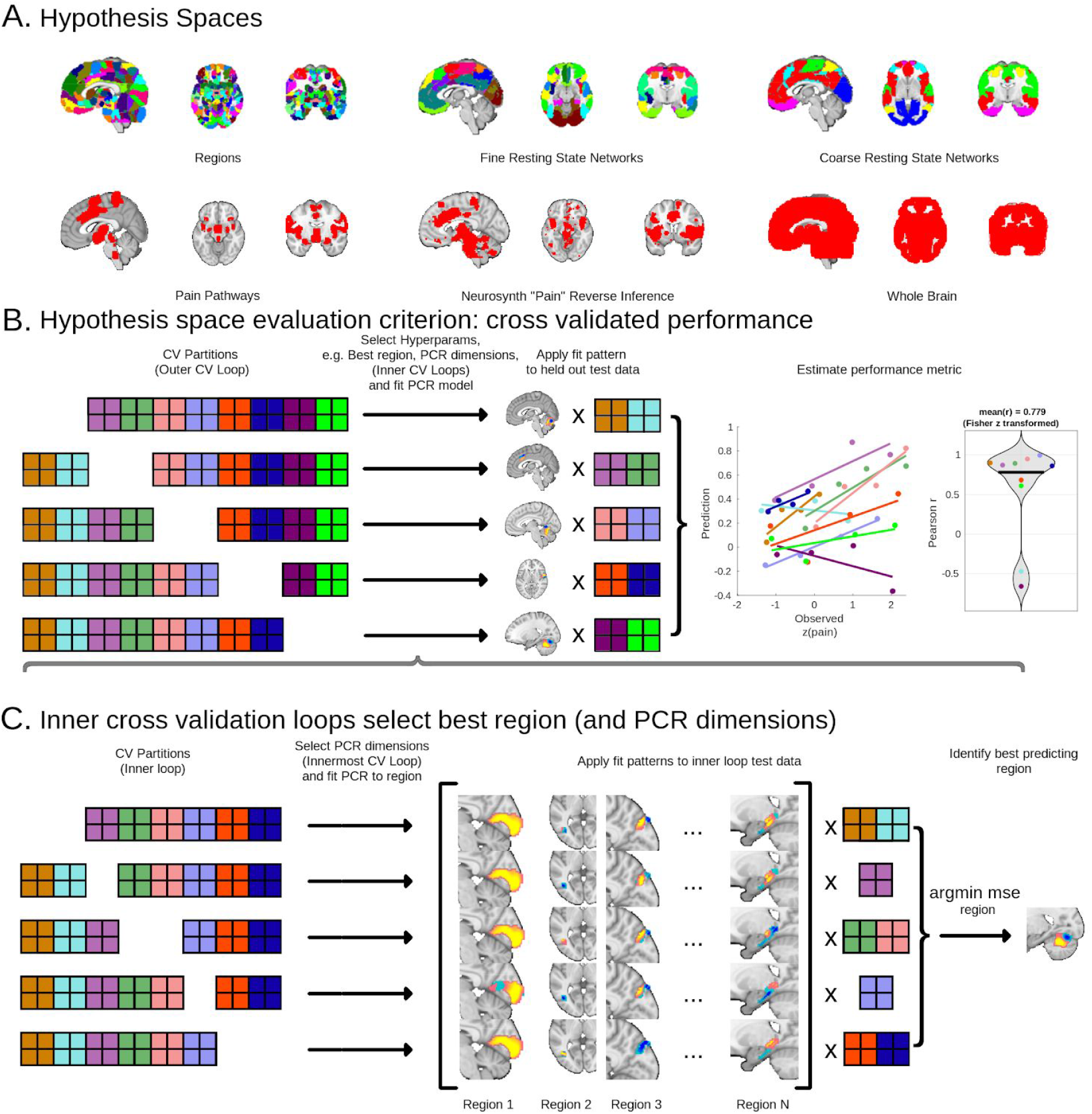
We test six different hypotheses regarding the scope of pain representation using cross validated principal component regression (PCR). (A) Three hypotheses (top row) test whether the BOLD signal of a single functionally homogeneous area best represents pain, and propose three homogenous area parcellations. Three additional hypotheses (bottom row) contend the BOLD signal of a set of distributed and functionally heterogeneous areas best represents pain. Proposed areas are illustrated. (B) We test each hypothesis based on cross validated MVPA predictions of pain intensity. We use principal component regression (PCR) to learn a model from hypothesized areas. We estimate model performance using 5-fold cross validation, training models on data averaged within pain intensity quartiles (smallest squares) for each subject (4x squares and one color per subject). Subject data is not fragmented across folds (shown), and studies are balanced across folds (not shown). We estimate model performance using within subject Pearson correlation of predicted and observed ratings. (C) The model fitting algorithm selects optimal model hyperparameters to minimise mean squared error, which it estimates using nested cross validation folds. In the case of multiarea hypotheses (A, top row), the algorithm treats region selection as a hyperparameter, and identifies a single best region for each outer fold (folds illustrated in B). This region may differ across outer cross validation folds. Principal component regression dimensionality is optimized by estimating expected mean squared error in an additional innermost cross validation loop (3 levels of nested folds). In the case of distributed area hypotheses (A, bottom row), the algorithm only performs the latter step (2 levels of nested folds). PCR hyperparameter optimization folds are not illustrated.

#### Elementary regions map

486 regions of interest subdivide the brain for anatomical characterization of sources of predictive information. These regions comprehensively cover brain gray matter from the brainstem through the forebrain, and are based on parcels and subcortical nuclei defined in previous published studies. We obtain 360 cortical regions (180 areas x 2 hemispheres) from the Human Connectome Project’s cortical parcellation, which delineates areas according to neuroimaging derived anatomical and multitask functional criteria (Glasser et al. 2016), transformed to volumetric MNI space. T1 and T2 based anatomical criteria delineate striatum, ventral tegmental area, substantia nigra, bed nucleus of the stria terminalis, pallidum, mammillary nuclei and the habenula (25 regions) based on (Pauli et al. 2016). T1 based anatomical criteria delineate the cerebellar lobules, vermis and deep nuclei (denate, interposed, fastigial; 34 regions total) (Diedrichsen et al. 2009). T1 derived tissue boundaries and white matter tractography delineate 16 thalamic nuclei and the hypothalamus (Krauth et al. 2010). A variety of nucleus specific functional and anatomical criteria define monoaminergic brainstem nuclei, the rostroventral medulla, parabrachial nuclei, trigeminal nuclei, nucleus ambiguus, nucleus tractus solitarius, tegmental nuclei, and some subdivisions thereof (24 areas) (Zambreanu et al. 2005; Fairhurst et al. 2007; Nash et al. 2009; Sclocco et al. 2016; Pauli, Nili, and Tyszka 2018; Shen et al. 2013; Keuken et al. 2014; Keren et al. 2009; Beliveau et al. 2015; Bär et al. 2016). Functional homogeneity criteria from a graph theoretic analysis of functional connectivity delineates remaining gross divisions of the brainstem (17 areas) (Shen et al. 2013). Finally, anatomical criteria adapted from post-mortem studies and included in the SPM Anatomy Toolbox subdivide hippocampal formation into CA1, CA2, CA3, dentate gyrus and subiculum, as well as gross subdivisions of the amygdala (superficial, latero-basal, centro-medial) (9 regions) (Amunts et al. 2005; Eickhoff et al. 2005).

#### Network parcellation maps

We used two different resting state network parcellations of the cortex as source regions for pain intensity prediction. One involves 7 networks (coarse resting state networks, cRSN), and the other is a finer parcellation of 17 networks (fine resting state networks, fRSN). cRSN regions are bilateral, while the fRSN regions are largely unilateral (32 regions total), and are derived in the same study sample (Yeo et al. 2011).

#### Pain Pathways Map

We manually create a map by combining areas from the elementary regions map (above) by selecting regions related to pain in previous literature (Apkarian et al. 2005; Duerden and Albanese 2013) and regions that play important roles in non-human pain studies. The areas included are ventral posterolateral, intralaminar and mediodorsal thalamic nuclei, hypothalamus, parabrachial nuclei, periaqueductal gray matter (PAG), rostroventral medulla, amygdala, dorsal posterior, middle and anterior insula, primary and secondary somatosensory cortices, and anterior medial cingulate cortex/medial prefrontal cortex (PFC). Regions were selected based on the prior knowledge of the field (by T.W.D.). The map and labels are available online in the Cognitive and Affective Neuroscience Laboratory Github repository at https://github.com/canlab/Neuroimaging_Pattern_Masks/.

#### Neurosynth.org empirical occurrence map

Following the approach of (Wager et al. 2013), we define an empirical map delineating regions which occur in neuroimaging studies of pain by searching neurosynth.org (Yarkoni et al. 2011) for “pain”, and thresholded the “association test” map (a quas-“reverse inference” map derived from frequentist test results) at the false discovery rate (FDR) q < 0.05.

### Multivariate Pattern Analysis (MPVA): General Method Description

Multivariate patterns are fit to each hypothesis space using principal component regression (PCR) with bayesian optimization of mean squared error (MSE) used to select the PCR dimensions. The dimensionality search space is restricted to be between 1 PCA dimension and R PCA dimensions, where R is the rank of the training data. We preclude intercept only models (0 PCA dimensions) *a priori*. Finally, we used nested 5-fold cross validation to estimate model generalization performance: 5-fold inner CV loop (Figure 2C) are used exclusively for bayesian selection of optimal PCR dimension (which is part of the model fitting procedure), and these are nested within 5 outer CV loops ultimately used for evaluating model performance (Figure 2B). We estimate model performance using Pearson correlation between predicted and observed values computed within-subject. This means the scope of this study is limited to within-subject variance in pain rather than between individual differences. Subject-wise Pearson r-values are normalized by Fisher’s *z* transformation. We repeat this entire nested cross validation procedure twice to stabilize results with respect to fold slicing (2×5 repeated k-fold cross validation). Results shown are averaged over repetitions. PCR implementation is available at https://github.com/bogpetre/mlpcr0.

We employ MVPA in two methods: (1) in a mega-analysis which pools data across studies and yields a single model for each hypothesis space (and an associated performance estimate), and (2) in an analysis of the learning capacity of each hypothesis space, which trains a separate model for each study (with seven associated performance estimates for each hypothesis space, one for each study’s model). The former gives us the best performing models we can obtain given our large dataset, and tells us how well we can expect them to perform (“model generalization performance”). The latter estimates how much can be learned from more typically sized datasets and allows for inference to how much will be learned by new models trained in new studies (“learner generalization performance”). Each method is described below in turn.

Bayesian hyperparameter optimization samples from the hyperparameter space (PCR dimension) and estimates loss (MSE) at each sampled point using (5-fold) cross validation. Gaussian process regression estimates the posterior distribution over loss functions after each sampling using the MSE from all thus-far obtained sample estimates, stopping after 30 iterations (our implementation’s default stop condition). There are two major advantages to this approach over other methods like grid sampling. First, cross validation is subject to sampling variance (different fold slicing produces slightly different results). Gaussian process regression models this variance as noise and takes this noise into account when selecting global optima. Second, this approach estimates a loss function after each sampling iteration, allowing for intelligent selection of the next best point to sample (for instance sampling in the domain which maximally improves precision of posterior loss function estimates). We used the algorithm as implemented in the BayesianOptimization toolbox in Matlab (Snoek, Larochelle, and Adams 2012).

### MVPA mega-analysis and model generalization performance

We perform a MVPA mega-analysis of data aggregated from seven training studies using nested 2×5 repeated k-fold cross validation. We train separate MVPA models for each hypothesis space pooling data across 7 studies. All input spaces require estimation of a PCR dimension. We estimate these dimensions in a nested cross validation loop as described in the “General Method Description” above, however modular spaces (elementary regions, cRSN and fRSN) require the additional selection of a best module. We use an additional level of cross validation fold nesting for this and sample modules exhaustively (so counting the nested 5-fold cross validation for optimization of PCR dimension within each area, so this is ultimately a 3x nested cross validation).

The mega-analysis balances participants across studies. Our smallest study had 16 participants, resulting in random sampling of 16 subjects from each study, for a total of 112 subjects and 6668 stimulus trials. After quartiling data this results in model training over 448 observed points.

### MVPA Mega-analysis Brain Map Visualizations

This study uses bootstrap resampling to generate brain maps representative of the study population.

Bootstrap resampling identified statistically significant voxels (uncorrected, bootstrapped 95% CI does not include 0) for multisystem predictive models (neurosynth and pain pathways) and the full brain predictive model. Additionally, we used bootstrapping resampling to identify significant voxels for each region, cRSN and fRSN selected as optimal by any inner cross validation loop. For visualization we fit predictive maps to the entire dataset (balanced across studies and subjects but not resampled), and then only show voxels identified as statistically significant under the bootstrap analysis.

For any bootstrapped analysis it is problematic to identify the PCR hyperparameters, namely the dimensionality of the underlying PCA. We wish to identify the relationship between PCA dimensionality and the expected generalization performance of the obtained PCR models, a procedure which generally relies on some form of cross validation (see section Multivariate Pattern Analysis: General Method Description). However, cross validation is not valid on bootstrapped samples due to likely redundancy of some resampled observations in both training and test splits. A common approach is to estimate model error as a function of the hyperparameters in the original, non bootstrapped sample, and then reuse that point estimate of the optimum dimension for PCR in the bootstrapped samples. However, there is some uncertainty in the estimate of the optimal hyperparameter which our Bayesian hyperparameter optimization scheme allows us to readily integrate into bootstrapped analysis.

Bayesian optimization of hyperparameters estimates a loss function of the hyperparameters (e.g. *error = f(dimensions)*) using Gaussian process regression, which entails computing a posterior probability distribution over loss functions. This posterior models uncertainty in the model estimate. We compute this posterior using the full dataset, and draw an instance *f\** from the posterior distribution over *f*, which varies according to the gaussian process model’s estimate of uncertainty in *f*, compute the global minima of *f\**, and use that as the hyperparameter for the bootstrapped PCR. This method is novel, results in greater MVPA map variance across bootstrap iterations, and more conservative bootstrap correction than typical in other MVPA studies.

### Study-wise MVPA and learner generalization performance

We define the learner as the combination of algorithm (PCR) and the hypothesis space (regional, cRSN, fRSN, pain pathways, neurosynth or full brain). For learner performance estimates we do not aggregate data across studies, instead defining 7 non-overlapping datasets, one for each study (using the complete subject sample available in each case, in contradistinction with the mega-analysis where subjects were balanced across studies), and train separate patterns for each. This study-wise MVPA uses a total of 171 subjects and 9773 trials (16-30 subjects, 256-2592 trials per study MVPA, table 2). As for the mega-analysis, we average data within pain rating quartile within subject, and evaluate the performance of MVPA patterns using 2×5 repeated k-fold nested cross validation.

### Bayesian hypothesis tests

We assess support for a null hypothesis using Bayes factors. Bayes factor tests are goodness of fit ratios that compare how well one model specification vs. another accounts for data while taking into account all possible model parameter values and their likelihoods. In all cases where we test support for the null we first begin with a frequentist test which fails to reject the null, i.e. an effect of interest lacks statistical significance (p > 0.05). This constitutes the ‘alternative model’. Null models tested are all nested instances of the ‘alternative’ model lacking this parameter which fell short of significance in frequentist testing. Non significant parameters of interest in alternative models often have mixed sources of error (e.g. random noise and subject or study related random effects). In such cases we remove all factors associated with parameters of interest, including associated random effects, in null models. Following (Jeffreys 1998), this study considers a Bayes factor greater than 10 in favor of the null “strong” enough to confirm the null.

We implement Bayesian models using markov chain monte carlo (MCMC) sampling algorithms specified in the STAN programming language (Carpenter et al. 2017). The rstanarm package efficiently and reproducibly converts our linear mixed effects model specifications used for frequentist testing into STAN source code for null hypothesis confirmation (Goodrich et al. 2018). We subsequently perform Bayes factor tests using bridge sampling (Gronau, Singmann, and Wagenmakers 2017). We evaluate four MCMC chains with a burn in of 5000 samples and subsequent drawing of 5000 samples each, and visually confirm chain convergence by inspecting parameter trace plots and by confirming unitary Rhat statistics. Additionally, to confirm convergence between frequentist and bayesian model specifications we compare maximum a posteriori (MAP) parameter estimates in null and alternative models with estimates obtained using restricted expectation maximum likelihood as implemented by the frequentist mixed effects modeling LME4 R package (Bates et al. 2015). This ensures models used for null hypothesis confirmation primarily reflect the data rather than Bayesian model priors.

### Mediation analysis

Mediation analyses explore MVPA derived models and facilitate their interpretation. We use a pair of ordinary least squares (OLS) regression models to estimate regression coefficients. First we regress an MVPA model prediction (the mediator) on stimulus intensity and psychological manipulations (independent variables, model a) and second we regress pain report (the dependent variable) on the mediator and independent variables (model b). We refer to the coefficient relating the mediator to the independent variable as *α*, the coefficient relating the mediator to the dependent variable as *β*. The coefficients’ product (*α*\**β*) estimates the mediation effect. Because resultant distributions are often highly skewed, we use bias-corrected bootstrap sampling to estimate confidence intervals of this product (MacKinnon, Lockwood, and Williams 2004) using 5000 resamplings of subjects, at the two-tailed alpha 0.05 level. This implementation is identical to that used by Mplus 7.11, except we resample subjects (i.e. sets of observations) instead of individual observations in accordance with independence assumptions underlying bootstrap tests. The resulting estimates are slightly more conservative than equivalent parametric estimates. We also model subject fixed effects (one constant term per subject), but because these effects do not vary across bootstrap iterations (bootstrapping is performed between not within subjects), bootstrap distributions for these effects are not computed. Instead, we orthogonalize dependent and independent variables and mediators with respect to subject fixed effects (one constant term per subject) *a priori* before estimation of all path coefficients. Thus, all mediation models reflect mediation of within-subject pain variance.

We select independent variables for mediation models in two steps. First, we consider sensitization and habituation effects as well as all experimental manipulations in a multivariate regression analysis on pain report. Second, backwards stepwise regression iteratively removes variables that do not significantly predict pain report. We evaluate all remaining variables for mediation. MVPA model predictions are used as mediators, but are not included in the variable selection process just described.

An example of our path models can be succinctly expressed in Wilkinson notation (Wilkinson and Rogers 1973) (common to linear model specifications in matlab and R), bearing in mind that subject intercepts are independently estimated as fixed effects here, as follows,
model a:

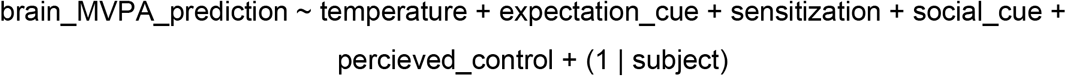

model b:

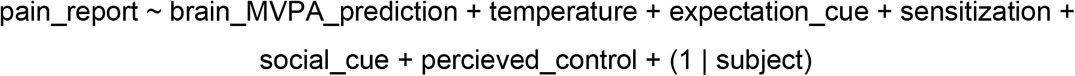

In this example, temperature, expectation cues, sensitization/habituation effects (note that sensitization also encompases habituation since the two only differ in sign), social cues and perceived control would all show statistically significant prediction of pain report in a multiple regression.

As for our behavioral analysis (see “Behavioral Analysis” above), we quantified mediation variance explained using the method of (Scherrer 1984). Whereas in the behavioral analysis we used standardized regression coefficients to scale the covariance of predicted outcome with independent variables, here we scale by the standardized indirect path coefficients (i.e. product of standardized *α* and standardized *β*, the coefficients of the indirect paths from models a and b, respectively). This value is equivalent to the marginal reduction in variance explained by mediated independent variables due to inclusion of the mediator (brain prediction) in the path b model, i.e. the reduction in variance explained by the corresponding direct effect on pain report.

## RESULTS

### Characterization of factors affecting pain

Behavioral models of pain report show different experimental factors drive pain across studies. All but one study (Study 6) have manipulations of thermal stimulus intensity across various ranges of intensity and duration (41.1-50C, 1-11s), and all but one study (Study 5) have psychological manipulations of pain (pain predictive expectation cues, Studies 1-4; social cues, Study 7; romantic partner handholding, Study 6; perceived control, Study 4). Variance in these experimental manipulations explain 51% of variance in pain report in a balanced aggregate of data across studies (n = 112 subjects, Figure 1B, Table 3), and 71% of within-subject pain variance, however this varies substantially between studies, ranging from primarily cognitive factors driving pain variation (Study 2) to primarily sensory factors like thermal stimulus intensity (Study 5). This reflects systematic differences in thermal stimulus intensity and duration as well as variation in efficacy of non-noxious manipulations between studies. There are also endogenous non-experimental factors, namely sensitization and habituation effects (Table 3, sens./habit.) or subject specific idiosyncrasies, which drive pain report and in some cases dominate pain report variance (Study 6). Differences are expected in sensitization and habituation effects based on the number of thermal stimuli and stimulation sites used (Jepma, Jones, and Wager 2014), while systematic differences in pain report across subjects will depend on the particular rating scales, calibration procedures and instructions provided in each study. Thus, variation in factors affecting pain across studies are consistent with underlying variations in study designs, and demonstrate how heterogeneous these data are.

### Characterization of MVPA models

Multisystem and whole brain hypothesis spaces are fixed, but modular hypothesis spaces are flexible: any particular area might be selected to predict pain. Indeed, many modules (51/486 elementary regions, 7/32 fRSN and 4/7 cRSN) are individually predictive (p < 0.05 regression coefficient, cross validated predicted vs. observed pain, random subject and study slopes and intercepts, Šidák correction for multiple comparisons, Figure 3), and no single region stands out as most predictive, with many regional predictions showing high sampling variance, suggesting the “best” region or network will highly depend on the subject sample selected (median sem, region: 0.10, fRSN: 0.10, cRSN: 0.10, z-Fisher transformed Pearson r across subjects, no random effects, n = 112, illustrated by whiskers in Figure 3). However, an examination of the distribution of predictive areas suggests certain trends might characterize “best” module selection. The areas which show high predictive accuracy, while numerous, are also systematic in their distribution in the brain, preferentially covering clusters of brain areas that are implicated in sensory processing, motor control, orienting attention to salient stimuli, ascending nociceptive targets in the midbrain like the periaqueductal gray (PAG) and parabrachial pigmented area (PMP, a subdivision of ventral tegmental area), and cortical nociceptive relays like ventral posterior medial thalamus (VPM). Additionally, predictive regions are disproportionately in the right hemisphere, reflecting the preferential contralateral representation of thermal stimuli delivered to the left arm (32/51 elementary regions and 5/7 fRSN right lateralized, one elementary region and all cRSN are bilateral). Consistent with this analysis we find high variability in module selection across cross validation folds (Figure 4), and regions selected appear to systematically sample from areas implicated in sensory processing and orienting responses, are overrepresented in the right hemisphere, and show some overlap in regions which are selected across modular spaces, with ACC regions regions appearing across hypothesis spaces, and areas which include the insula being selected in RSN spaces (Figure 3).

**Figure 3.**
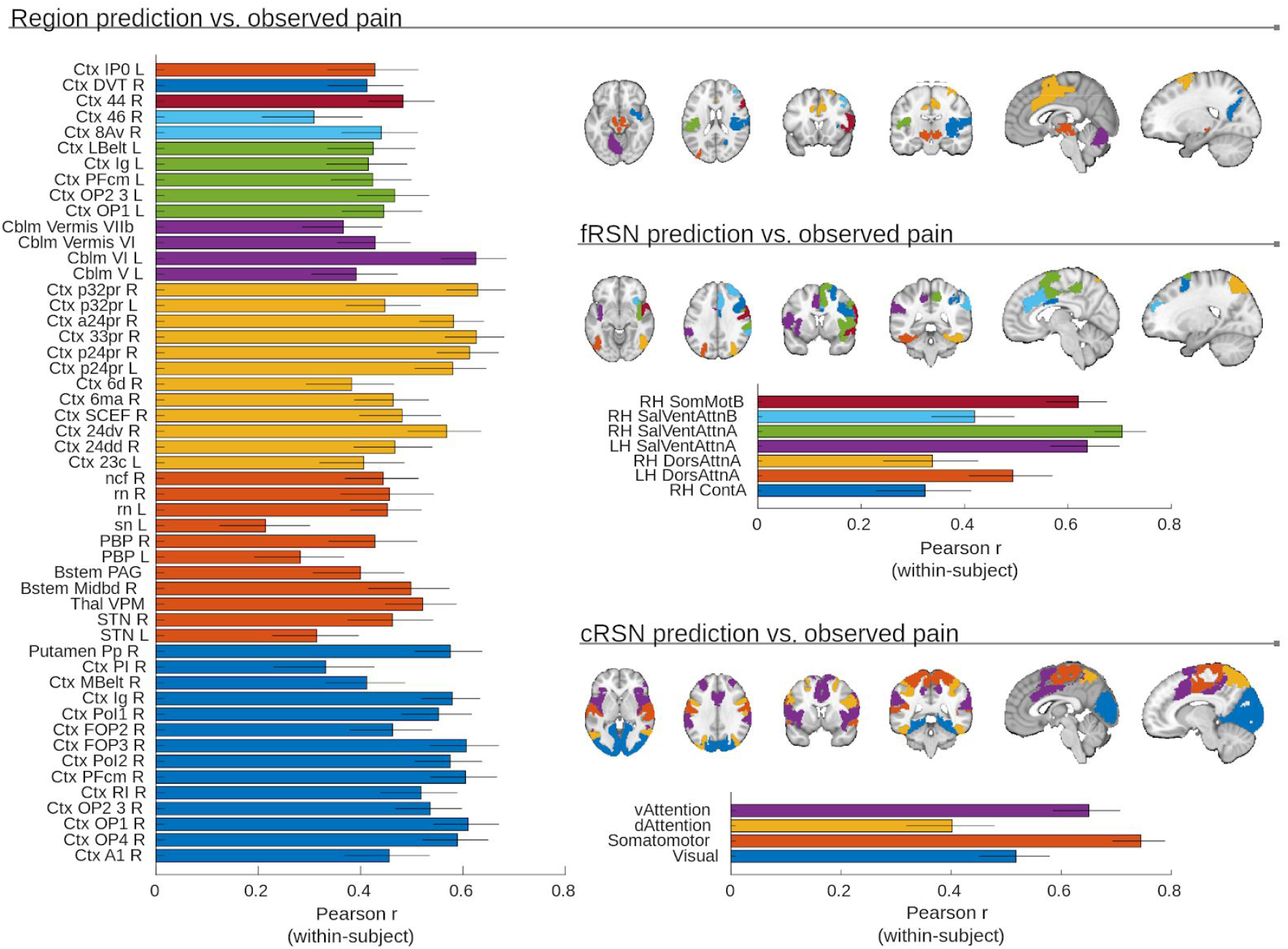
Numerous brain areas yield models with significant pain decoding capabilities, but prediction accuracy varies considerably across participants (s.e.m. whiskers shown, n = 112). Predictions from 51 elementary regions, 7 fRSN and 4 cRSN correlate significantly with pain report (shown; *ɑ* = 0.05, twice repeated 5-fold cross validated predictions modeled with random subject and random study effects, Šidák correction for multiple comparisons, effective p < 10e-4). This means the “best” region, cRSN or fRSN will be highly dependent on the subject sample examined (and will differ across folds in a cross validation scheme). Regions color coded by contiguous areas. cRSN and fRSN colored by individual network parcels.

**Figure 4.**
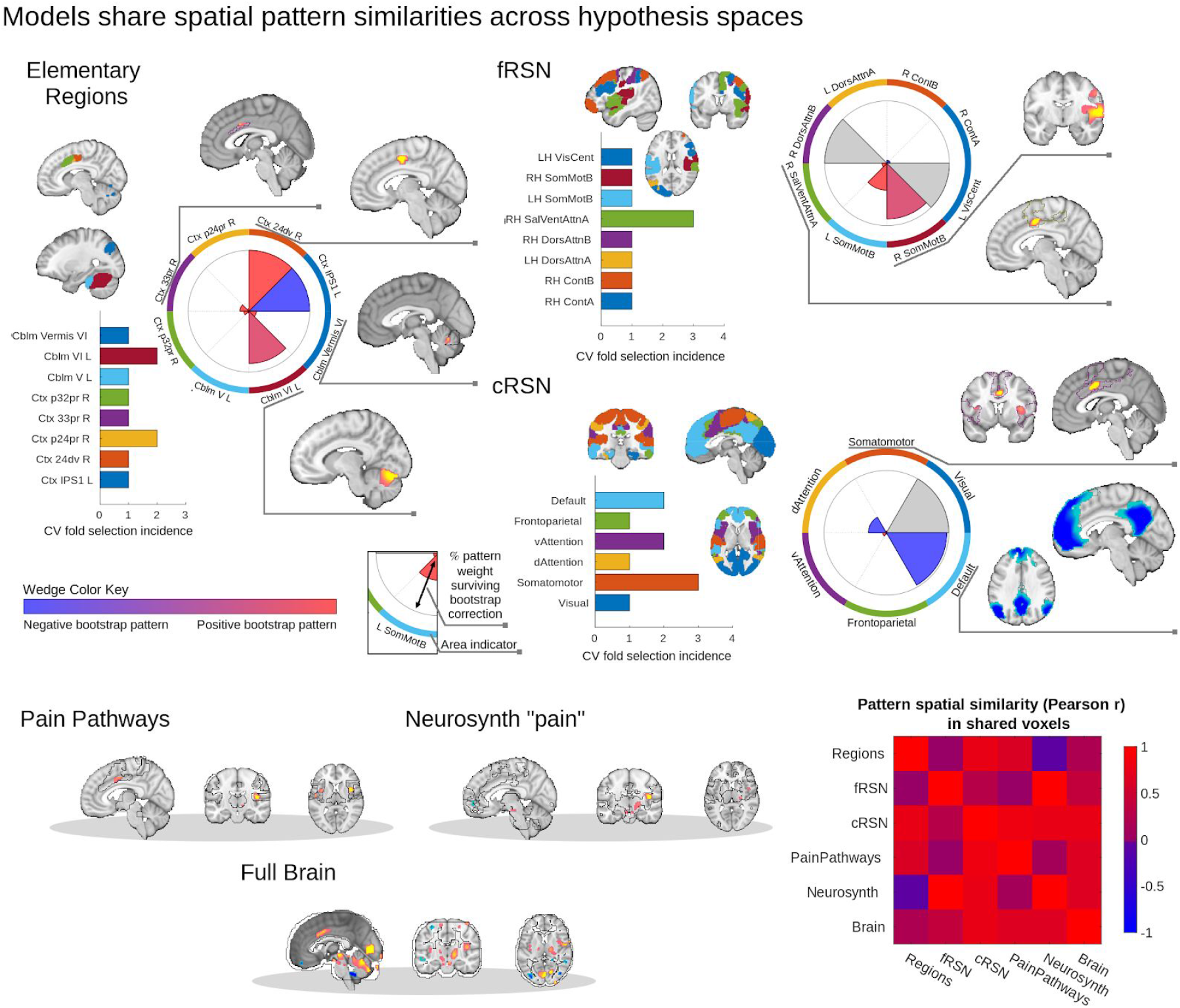
Machine learning yields brain models of pain response which can be interpreted as nested models which draw on increasing amounts of unique information. Modular hypotheses (regional, fRSN and cRSN) vary in region selection across folds. Bar graphs illustrate area selection incidence, which is highly variable across folds, consistent with a large number of highly predictive regions and network parcels. Bootstrap corrected maps are computed for all regions to characterize pattern stability across folds, since patterns will vary even if different folds select the same region (although most modules are only selected once). Representative maps are shown. Wedge plots summarize maps in terms of net pattern weight of bootstrap corrected maps (red: positive vs. blue: negative) and percent of pattern that survives bootstrap correction (wedge radius; gray: highly unstable patterns, no significant bootstrap voxels). Multisystem and full brain patterns are summarized only in terms of bootstrap corrected maps. While the relatively low number of significant weights suggests low model stability with respect to subject samplings, a pattern spatial similarity matrix illustrates the relatively high stability of weights across model families (only conjunction of bootstrap corrected voxels considered). Across all six model families we commonly observe frontal midline deactivations, and cingulate, insular and cerebellar activations, supporting a nested hypothesis space interpretation. 5008 bootstraps, voxel-wise p < 0.05, uncorrected. Outlines illustrate unthresholded model boundaries.

Pain pathways, neurosynth and full brain models cannot vary in voxel selection across folds the way region or network patterns do (each fold fits weights to the same voxels), however each fold still differs in configurations of MVPA patterns. Pattern stability is most succinctly illustrated by voxel maps of pattern weights. Bootstrapped distributions of pattern weights across subjects show stable patterns in ACC, insular, and visual cortices, PAG, cuneiform, parabrachial, and raphe nuclei of the brainstem, select cerebellar lobules, and pulvinar, VPM, and ventrolateral nuclei of the thalamus (significant percent voxel coverage in constituent regions relative to permuted null distribution, α = 0.05 uncorrected p < 0.001).

Patterns which correlate with pain in one hypothesis space remain correlated with pain in other hypothesis spaces, and conversely those that are anti-correlated with pain in one hypothesis space are also anti-correlated in models obtained in other hypothesis spaces (mean Pearson r: 0.48, interquartile range: [0.11, 0.81], 15 unique pairwise comparisons). Thus, the result of full brain hypothesis space stability analysis largely summarizes the most frequent patterns selected across spaces (Figure 4). For instance, areas like ACC, insula and cerebellum with significant positive weights in the pain pathways or neurosynth derived patterns also have significant positive weights in the full brain pattern. Similarly, negative pattern weights in the ventromedial PFC and dorsal attention areas of the full brain model (significant, α = 0.05) are reproduced by patterns fit in analogous areas in the fRSN and cRSN parcels. This suggests that as we move from spatially constrained models to spatially less constrained models we are in fact iteratively building on prior models in a consistent way. Larger models are leveraging the same signals as smaller models and are using them in a similar way, but also incorporate some marginal additional information available due to the broader scope of a larger hypothesis space. Thus, each model draws on progressively greater amounts of independent additive information across brain areas as we move from elementary regions to network parcellations to multisystem and full brain patterns.

### MVPA mega-analysis and model performances

Mega-analytic comparison of predictive performance across models (n = 112 subjects, 16 per study) show significant differences in model performance (F_5,11_= 11.12, p = 0.021, mixed model, random subject and study intercepts, random study effects, Satterthwaite correction for degrees of freedom (df); Figure 5A, Tables 4, 5), and the more cortical areas a model aggregates the better its performance. Planned comparisons show models from multisystem (*a priori* “pain pathways” and empirical, neurosynth derived) and full brain spaces are more predictive on average than modular patterns (regional, fRSN and cRSN parcellations; F_1,6.9_ = 14.9, p = 0.006). Subdividing further, signatures derived from modular hypothesis spaces perform similarly to one another on the one hand and on the other hand models derived from multisystem or full brain spaces show equivalent performance with one another (mixed effects contrast not significant, Bayes Factor in favor of the null, cRSN vs. fRSN: 14.0, roi vs. c/fRSN: 98.3, neurosynth vs. pain pathways: 232.3, full brain vs. neurosynth/pain pathways: 42.0). However, a statistical contrast of multisystem and full brain spaces with modular spaces implicitly performs model averaging (average of pain pathways, neurosynth and full brain on the one hand vs. cRSN, fRSN and elementary regions on the other), and this complicates interpretation. For greater transparency and interpretive convenience we also perform a post-hoc comparison of the specific difference between full brain pattern performance (our largest model) and the elementary region’s performance (our smallest model), and find the full brain pattern is significantly more predictive (F_1,7.9_ = 12.4, p = 0.008). This demonstrates the added brain tissue aggregated under the pattern weights of our larger models carry unique pain related signals that cannot be subsumed under more limited subsets of brain areas.

**Figure 5.**
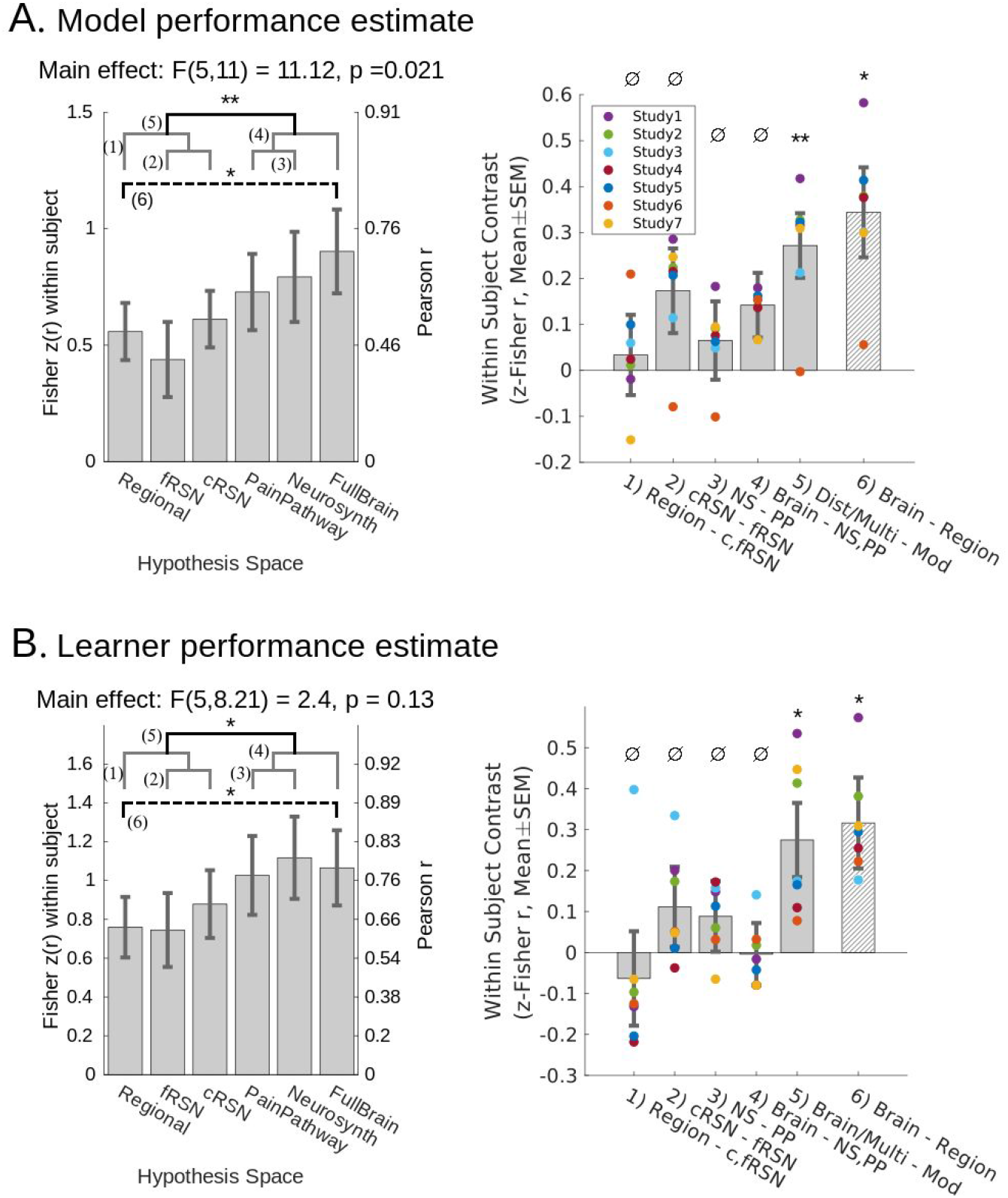
Multisystem and full brain representations of pain perform better than modular representations of pain in a 2×5 repeated cross validation test of generalization performance. We Fisher z-transform within subject correlation coefficient (predicted vs. observed) to quantify performance. (A) All obtained models show significant predictive performance (p < 0.004, random subject and study intercept, random study effect), and the specific models obtained here to predict pain using distributed sets of brain areas will outperform the models trained on individual modules (planned comparison, mean(PainPathway, Neurosynth, FullBrain) - mean(regions, fRSN, cRSN) = 0.272(0.07), F_1, 6.9_ = 14.9, p = 0.0064). We confirm the nullity of remaining planned contrasts (Bayes factor, BF > 10 favoring the null). Post-hoc comparison shows a full brain model outperforms the best single region (FullBrain – Regions = 0.34(0.10), F_1,7.9_ = 12.35, p = 0.008). Models are trained after pooling data across seven datasets, yielding 90 subjects per training fold. (B) We retrain cross-validated models separately for each study to infer whether findings generalize to new PCR models trained on new datasets. All obtained models show significant predictive performance (p < 0.001, random subject and study intercept, random study effect), but comparison of performance across models from each study shows that distributed representations will continue to outperform modular representations (planned comparison, mean(PainPathway, Neurosynth, FullBrain) - mean(regions, fRSN, cRSN) = 0.27(0.09), F_1,6.1_ = 9.04, p = 0.022). Remaining planned contrasts are null (BF > 10). Post-hoc comparison shows training on the full brain will result in better performance than training on elementary regions (FullBrain – Regions = 0.32(0.11), F_1,5.2_ = 7.96, p = 0.034). Training set sizes = {22, 14, 13, 23, 21, 24, 20} subjects, resp. All effects estimated with subject-wise repeated measures and random study effects. Mean within study effects overlaid as colored dots. Gray brackets: nonsignificant planned comparisons; black bracket: significant planned comparison; dashed bracket: significant post-hoc comparison. Contrasts enumerated across brackets. *p < 0.05, *p < 0.01, ^∅^confirmed null. Mixed effect degrees of freedom estimated using Satterthwaite’s approximation. s.e.m error bars.

**Table 4.** Model and learner performance across hypothesis spaces (figure 5a-b)

|  | Model performance |  | Learner performance |  |
| --- | --- | --- | --- | --- |
| Hypothesis Space | r-value<sup>†</sup> [CI<sub>0.95</sub>] | df | r-value<sup>†</sup> [CI<sub>0.95</sub>] | df |
| Region | 0.507<br>[0.31, 0.67] | 6.58 | 0.636<br>[0.37, 0.81] | 6.42 |
| fRSN | 0.413<br>[0.18, 0.60] | 6.70 | 0.641<br>[0.28, 0.84] | 6.01 |
| cRSN | 0.545<br>[0.24, 0.76] | 6.24 | 0.701<br>[0.82, 0.86] | 6.23 |
| Pain Pathways | 0.623<br>[0.34, 0.80] | 6.17 | 0.773<br>[0.49, 0.91] | 6.12 |
| Neurosynth | 0.661<br>[0.31, 0.85] | 6.08 | 0.806<br>[0.54, 0.93] | 6.18 |
| Full Brain | 0.718<br>[0.43, 0.87] | 6.13 | 0.788<br>[0.54, 0.91] | 6.25 |

<sup>†</sup>Cross validated model performance (Pearson-r) within-subject is normalized by z-fisher transformation and modeled using a mixed effects design (random subject intercept, random study intercept and slope). Inverse z-Fisher transformed model coefficients and CI shown. CI<sub>0.95</sub>, 95% Confidence Intervals. df, degrees of freedom estimated using Satterthwaite's correction.

**Table 5.** Contrasts of hypotheses spaces across model and learner performances (figure 5a-b)

| Contrast<sup>†</sup> | Model performance contrast |  |  | Learner performance contrast |  |  |
| --- | --- | --- | --- | --- | --- | --- |
|  | z(r)<sup>‡</sup><br>(s.e.m.) | df | p-value | z(r)<sup>‡</sup><br>(s.e.m.) | df | p-value |
| multisys - local | 0.272<br>(0.07) | 6.88 | 0.006* | 0.275<br>(0.09) | 6.13 | 0.022* |
| brain - (NS & PP) | 0.141<br>(0.07) | 18.63 | 0.059 | -0.004<br>(0.08) | 9.79 | 0.959 |
| NS - PP | 0.065<br>(0.09) | 13.36 | 0.459 | 0.088<br>(0.09) | 13.22 | 0.325 |
| cRSN - fRSN | 0.173<br>(0.09) | 10.51 | 0.088 | 0.111<br>(0.10) | 6.86 | 0.293 |
| region - c/fRSN | 0.034<br>(0.09) | 6.45 | 0.713 | -0.063<br>(0.12) | 6.17 | 0.607 |
| brain - region (post-hoc) | 0.344<br>(0.10) | 7.92 | 0.008* | 0.316<br>(0.11) | 5.23 | 0.034* |

<sup>†</sup>Performance contrasts are modeled using a mixed effects design (random subject intercept, random study intercept and slope). <sup>‡</sup>z-fisher transformation of cross validated model performance (within-subject Pearson-r). df, degrees of freedom estimated using Satterthwaite's correction. NS, neurosynth; PP, pain pathways. \*p < 0.05

Regression of pain report on experimental manipulations shows pain signals originate from multiple sources (Figure 1B), including temperature, expectation, sensitization/habituation, social cues and perceived control. Additionally, experimental manipulations and our best brain models predict similar amounts of pain variance (70% and 72%, respectively), suggesting brain models may reflect the effect of experimental manipulations, so we perform a within-subject mediation analysis to evaluate which of these pain sources are captured by our best model (Figure 6, Table 6). The full brain model significantly mediates the effects of temperature (T_ind_: 0.065 [0.031, 0.113], standardized *α*\**β* [95% confidence interval], 12% mediated), expectation (exp_ind_: 0.030 [0.010, 0.060]), 12% mediated), and habituation effects on pain report (habit_ind:_ −0.022 [−0.051, −0.004], 10% mediated). However the full brain model does not fully capture these effects since direct effects remain significant in all cases (T: 0.471 [0.388, 0.577], partial-r^2^ = 0.34; exp: 0.227 [0.148, 0.302], partial-r^2^ = 0.14; social: 0.212 [0.177, 0.268], partial-r^2^ = 0.06; habituation: −0.139 [−0.211, −0.072], partial-r^2^ = 0.08; control: −0.073 [−0.162, −0.001], partial-r^2^ = 0.01; brain: 0.189 [0.107, 0.276], partial-r^2^ = 0.1). Brain model predictions mediate the effects of experimental manipulations because predictions share variance with the manipulations (brain partial-r_mediation_^2^ = 0.07, Figure 6 “*α*\**β*” inset), but brain model predictions also contain additional unique information (brain partial-r_unique_^2^ = 0.03, ‘model b’ total brain partial-r^2^ = 0.10). This shows our full brain model captures multiple dimensions of pain report, in proportions representative of their overall effect on experimental pain, but misses the effects of some variables (social cue and perceived control) while predicting something which cannot be explained by experimental manipulations alone.

**Figure 6.**
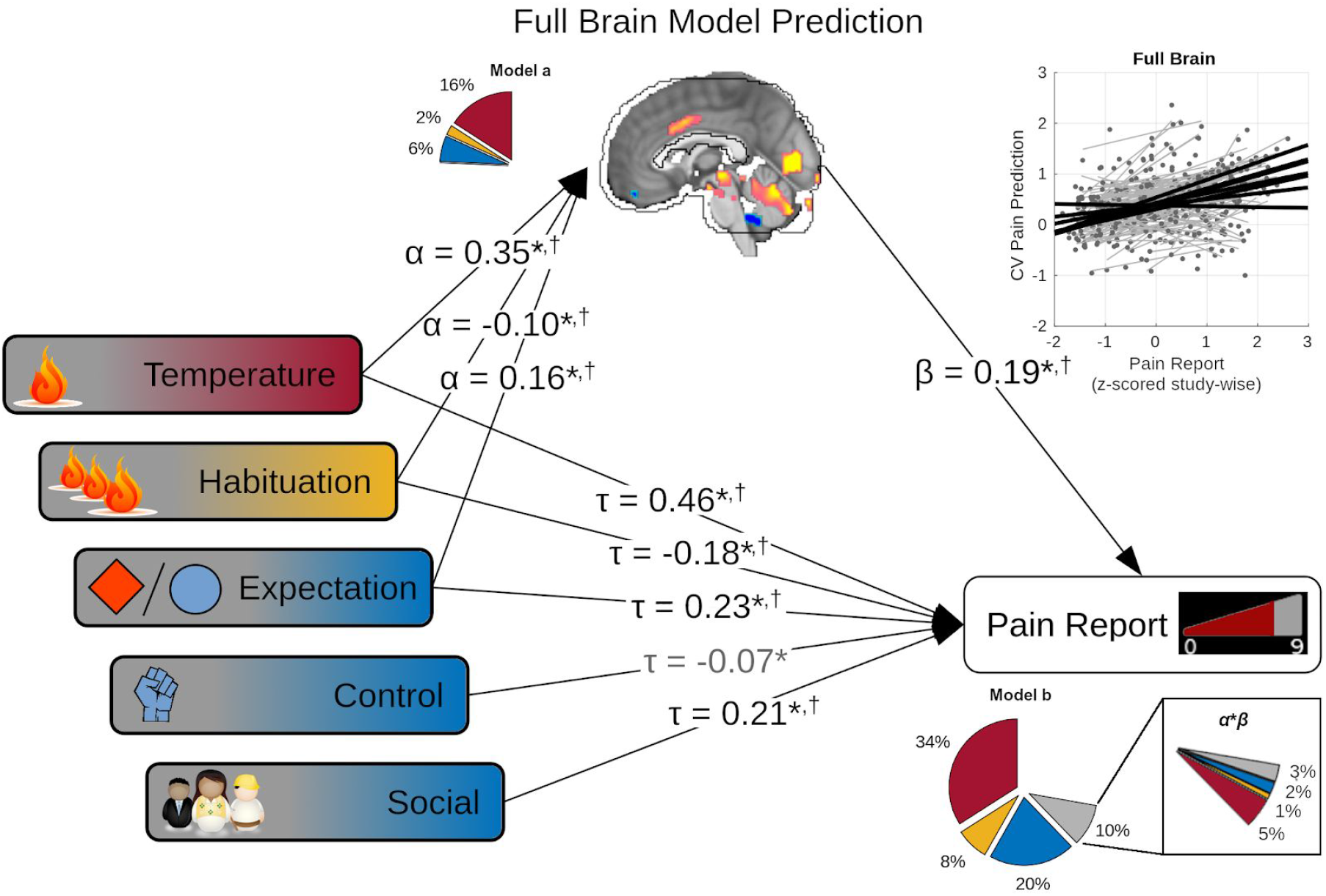
Multisystem models capture brain correlates of both sensory (red, orange) and cognitive (blue) influences on pain report. Five experimental factors show significant effects on pain report: temperature, habituation, perceived control (control), expectation and social cues (multivariate regression, fixed subject effects, p < 0.05). However, the cross validated (CV) full brian model predictions also significantly correlate with pain outcome (scatter plot), and temperature, habituation and expectation in turn significantly correlate with these full brain model predictions (*α*, standardized multivariate regression coefficients; Model a pie chart show partial-r^2^, color coded by factor). Finally, the full brain model continues to significantly predict pain outcome (*β*) even while controlling for experimental factors (*τ*). This suggests temperature, habituation and expectation effects are captured by the full brain model. Multivariate mediation analysis supports this explanation showing both significant indirect (mediated, *α*\**β*), and direct (*τ*) effects on pain report, with additional significant direct effects from perceived control and social cues. Together, these factors and the full brain model predictions explain 72% of pain variance (Model b r^2^, subdivided by color coded categories in pie chart; gray: brain model predictions). Of the 10% of variance explained apportioned to the brain predictions in multivariate analysis (model b pie chart), 7% represents mediated temperature, expectation and habituation effects (*α*\**β* pie chart insert, subdivisions rounded to nearest percent), however brain model predictions also explain some unique variance (residual gray wedge in insert). Together these results show multisystem brain models capture a multidimensional pain experience. Nearly identical conclusions are supported by full brain learner predictions (^†^, table 6). Standardized path coefficients shown. All parameter estimates computed after removing subject fixed effects. *p < 0.05, in models using full brain model predictions (shown). ^†^p < 0.05 in models using full brain learner predictions. Bias corrected bootstrap test for non-zero *α*, *β*, *τ*, and *α*\**β*. For details refer to table 6.

**Table 6.** Mediation model (standardized) coefficients and 95% confidence intervals (Figure 6)

| Factor | Using Full Brain Model Prediction |  |  | Using Full Brain Learner Prediction |  |  |
| --- | --- | --- | --- | --- | --- | --- |
| | $\alpha$ | $\beta$ | $\alpha*\beta$ | $\alpha$ | $\beta$ | $\alpha*\beta$ |
| Temperature | 0.346*<br>[.21, .49] | 0.463*<br>[.38, .57] | 0.065*<br>[.03, .11] | 0.508†<br>[.40, .61] | 0.429†<br>[.35, .51] | 0.129†<br>[.09, .17] |
| Habituation | -0.10*<br>[-.21, -.00] | -0.182*<br>[-.26, -.11] | -0.019*<br>[-.05, -.00] | -0.078†<br>[-.16, -.01] | -0.149†<br>[-.21, -.09] | -0.020†<br>[-.04, -.00] |
| Expectation | 0.16*<br>[.05, .27] | 0.226*<br>[.15, .30] | 0.030*<br>[.01, .06] | 0.111†<br>[.01, .21] | 0.192†<br>[.12, .26] | 0.028†<br>[.00, .05] |
| Control | 0.042<br>[-.09, .14] | -0.073*<br>[-.16, -.01] | 0.008<br>[-.02, .03] | -0.007<br>[-.10, .07] | -0.060<br>[-.13, .01] | -0.002<br>[-.03, .02] |
| Social | 0.04<br>[-.08, .07] | 0.208*<br>[.17, .26] | 0.000<br>[-.01, .02] | 0.037<br>[-.01, .09] | 0.221†<br>[.19, .25] | 0.009<br>[-.00, .03] |
| Brain | - | 0.187*<br>[.11, .27] | - | - | 0.254†<br>[.19, .32] | - |

\*†p < 0.05, bias-corrected bootstrap confidence interval, 5000 repetitions. Factors included based on statistical significance in multiple regression on pain (t-test, p < 0.05). All parameters and statistics estimated while modeling subject fixed effects (not shown).

### Learner generalization performance

Although the above mega-analysis and associated mediation analysis provides an estimate of expected predictive performance for our models in novel data, and informs the sources of information these models capture, it does not show if multisystem and full brain hypothesis spaces will systematically confer an advantage in pattern training in general. To establish whether multisystem and full brain patterns in general (rather than the specific patterns obtained here) outperform modular models developed in new datasets, we evaluate learner performance across individual studies (n = 16-30 subjects each, 7 studies, 171 subjects total). Regardless of a study’s design or experimental manipulation there is a systematic tendency for patterns trained on multisystem and full brain hypothesis spaces to outperform patterns trained on modular hypothesis spaces (F_1,6.1_ = 9.04, p = 0.022, mixed model, random subject and study intercept, random study effects, Satterthwaite df correction; Figure 5B, Tables 4, 5). It is important to stress that this is a stronger claim than the results of the mega-analysis since it shows more will be learned in a formal sense by allowing pain representation to span multiple brain systems than by adopting a localized perspective, and we quantify precisely how much more (Tables 4, 5). For instance, the best region approach only explains 81% of the signal represented by the full brain (ratio of r^2^).

Mediation analysis of study specific models does not show any significant mediation effects (Study 7 social mediation p < 0.1, Study 2 temperature mediation p < 0.1, all others p > 0.1), however this might be due to lack of power in small samples, so we pool each prediction obtained from each study’s full brain model and perform a within-subject mediation analysis across studies (Table 6). Backwards stepwise regression of all experimental manipulations shows significant effects for temperature, expectation, social cue, perceived control and sensitization/habituation, so all are evaluated for mediation by the full brain models’ predictions. Temperature and expectation show significant partial mediation (T_ind_: 0.129 [0.088, 0.175], 23% mediated; exp_ind_: 0.028 [0.005, 0.056], 13% mediated), while temperature, expectation, social cues, and habituation show significant direct effects on pain report (T_dir_: 0.429 [0.353, 0.510], partial-r^2^ = 0.31; exp_dir_: 0.192 [0.122, 0.259], partial-r^2^ = 0.10; social_dir_: 0.221 [0.188, 0.251], partial-r^2^ = 0.06; habit_dir_: −0.149 [−0.207, −0.092], partial-r^2^ = 0.06) while controlling for brain predictions (brain_dir_: 0.253 [0.186, 0.315], partial-r^2^: 0.16, partial-r^2^_mediation_: 0.12, partial-r^2^_unique_: 0.04). These results mirror those found in the mega-analysis (Figure 6, Table 6), albeit with perhaps a greater bias towards capturing pain over expectation or habituation effects. It may not be possible to understand which specific experimental factors are mediated by multivariate models in any particular study. Nor is it possible to distinguish whether any individual model captures multiple factors simultaneously, or if models obtained from different studies rather differ in the factors they mediate. However, these results do show that representations of multiple distinct dimensions of pain experience are learned across multiple applications of these models, meaning distinct sensory and cognitive pain modulating signals are successfully captured by multisystem spaces in a general sense.

### Validation

We validate both model and learner generalization performances by evaluating 3 studies which once again differ in experimental design, but which we have not used for model training or obtaining estimates of model or learner performances up to this point. Variability in model performances across validation studies fits estimates of variability in model performance well, suggesting our models are well calibrated (Figure 7A, 95% confidence interval (CI) estimates: gray violin plots, observed: connected dots). We also inspect significant planned and post-hoc comparisons from the model development stage in the validation studies, and find validation data falls within the range of expected effect sizes for multisystem and full brain vs. modular comparisons but not brain vs. elementary region comparisons. Multisystem and full brain models show significantly better predictions than modular models (F_1,575_ = 11.9, p = 0.0006, mixed model, fixed study effects, random subject intercept, Satterthwaite df correction), and while full brain predictions are not significantly different from elementary region predictions in the validation data (p = 0.68, BF = 3.3e4 in favor of null), in absolute terms full brain predictions still perform better than regional models on average across validation sample subjects. Both contrasts found to be significant for learner generalization performance and validation data fall within the expected range for all significant planned and post-hoc contrasts (multisystem and full brain vs. region, cRSN, fRSN: F_1,575_ = 38.2, p < 1e-8; brain vs. region: F_1,115_ = 5.6, p = 0.02; Figure 7B). These comparisons serve as a sanity check on the cross validated estimates of model and learner performance and confirm the validity of inferences drawn: in novel subjects and experimental designs multisystem and full brain models will systematically outperform more spatially constrained and targeted models of pain intensity.

**Figure 7.**
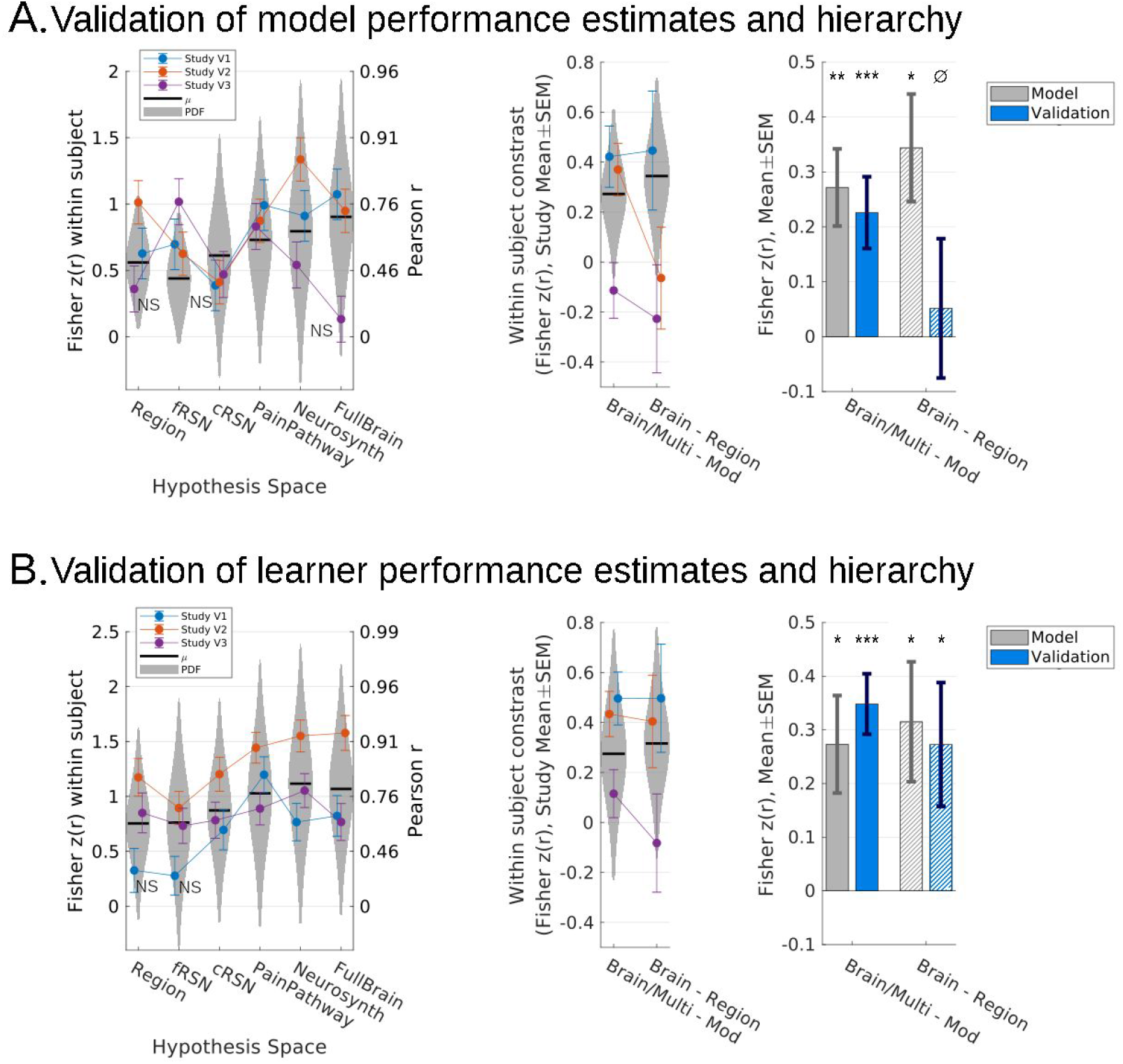
Distributed representations continue to outperform modular representations in validation datasets, consistent with obtained estimates of generalization performance. (A) All models fit to the 7 study dataset significantly predict pain in novel data, except for 3 models in 3/21 cases (p < 0.05, left, only non significant results labeled, Holm-Sidak corrected). Performance in validation data continues to be significantly better for multisystem and full models relative to modular models in validation studies 1 and 2 (mixed effects model, random subject intercept, F_1,160_ = 12.6, p < 10e-3; F_1,220_ = 11.8, p < 10e-3; resp.) as well as overall (mixed effects model, random subject intercept, fixed study effects, F_1,575_ = 11.9, p < 1e-3), however the full brain model does not show better performance than the best elementary region model specifically (BF = 3.3e4 in favor of null; right, only significant results indicated). (B) Nevertheless, retraining all six families of algorithms in novel data reproduces both findings that full brain and multisystem representations predict more pain than modular representations and the finding that a full brain model specifically will outperform any single best region. In a 2×5 repeated cross validation scheme all models separately retrained for each study successfully predict pain, except for 2/21, specifically in validation Study 1 (left, only non significant results labeled, Holm-Sidak corrected), however full brain and multisystem models significantly outperform modular models in validation Studies 1 and 2 (mixed effects, random subject intercept, F_1,160_ = 15.0, p = 0.0002; F_1,220_ = 28.4, p < 1e-6; resp.) and overall (mixed effects, random subject intercept, fixed study effects, F_1,575_ = 38.1, p < 1e-8). Additionally, the full brain models specifically outperform the best elementary region models in Study 2 (mixed effects, random subject intercept, F_1,44_ = 10.1, p = 0.0027) and overall (mixed effects, random subject intercept, fixed study effects, F_1,115_ = 5.59, p = 0.020; right, only significant results indicated). BF, Bayes factor. ^NS^p > 0.05 (not significant), *p < 0.05, **p < 0.01, ***p < 0.001, ^∅^BF > 10, “strong” evidence of null. Gray bars previously shown in figure 4. Error bars: SEM, random subject intercepts, random study (gray bars) or fixed study (otherwise).

## DISCUSSION

This study investigates whether pain representation is best characterized as a localized signal within a discrete module, a distributed signal across multiple brain systems, or if it is represented throughout the brain. In a mega-analytic sample of subjects pooled across seven studies we find multisystem and whole brain models best decode evoked pain. However, models trained on the full brain show no advantage over models trained on multisystem spaces, suggesting the latter have already subsumed all uniquely informative signals, while the elementary region account emerges as a surprisingly good approximation of overall pain representation. Followup comparisons of such models trained separately for each study support identical conclusions, showing these results will generalize to new models which might be learned from novel subjects, experimental designs and studies. These results are consistent with a tripartite theory of pain representation (R. Melzack and Casey 1968) which entails a pain “neuromatrix” distributed across sensory, limbic and prefrontal brain areas, but build on prior studies by formalizing both the scope of information representation and its configuration across a diverse collection of task conditions while generalizing to heterogeneous evoked pain experiences. We thus demonstrate evoked pain is represented preferentially by specific brain areas but is nevertheless better seen as spanning multiple brain systems.

Unlike traditional univariate approaches which focus on localizing brain activity in response to a stimulus, the multivariate predictive models we employ are formal models which map a specific configuration of brain activity to a construct of interest (Kragel et al. 2018). Multivariate modeling has been successfully applied to predict pain intensity, analgesic drug efficacy (Wager et al. 2013), pain reappraisal (Woo et al. 2015), and distinguish physical pain from social (Koban et al. 2019) and vicarious pain (Krishnan et al. 2016), often by making liberal judgments about the spatial extent of pain representation. We formally define spatial scales largely according to established biological gradients in anatomy and function (elementary regions, cRSN and fRSN, pain pathways), or a meta-analytic area representative of the neuroimaging findings of the field (neurosynth). Preliminary systematic attempts to investigate spatial scale have suggested pain is distributed throughout the brain (Kragel et al. 2018; Brodersen et al. 2012; Brown et al. 2011), but due to methodological and design limitations do not fairly compare across spatial scales, especially network or multisystem scales, or allow for inference to new individuals or diverse experimental conditions. Our application of MVPA across nested scales in a large dataset, spanning multiple experimental paradigms, allows us to characterize the architecture of evoked pain in a general sense.

Despite the strengths of multivariate brain models, this study moves beyond multivariate modeling to estimate the scope of information content. We operationalize “brain information content” as what can be learned in general rather than any particular instance of learning. Focusing on abstract information content benefits from the strengths of machine learning which is optimized to make accurate predictions by maximally exploiting available information even when the underlying generative processes cannot be recovered (Næs and Martens 1988; Geman, Bienenstock, and Doursat 1992). However, directly generalizing from a particular model’s performance to new models which might be learned is notoriously problematic (Bengio and Grandvalet 2004). Our mega-analysis yields specific multisystem models which outperform specific regional and network models when applied to new studies and subjects. Validating these models through direct application to novel datasets (without model retraining) illustrates the rigid scope of this and traditional MVPA approaches. We circumvent the problems of using a model instance to estimate the learning capacity of a class of models by simply examining performance of independent models retrained for each study separately (i.e. 42 different models, 6 hypotheses, 7 studies), following precedent established in machine learning (Dietterich 1998; Boulesteix et al. 2015), and applications in genomics (de Souza, de Carvalho, and Soares 2010). Variability in model performances across studies informs the performance of new unseen models trained on unseen new subjects in new studies, and quantifies the information content which can be recovered at each scale across experimental contexts in a practical sense. We successfully validate these estimates and illustrate the broader nature of this claim by training new models in new studies on new subjects and obtaining an identical performance hierarchy favoring multisystem models. This provides a blueprint for exploiting large heterogeneous datasets to practically quantify information content in the brain, one which can just as readily be applied to other brain phenomena besides pain.

Although our work primarily supports a multisystem representation of pain, local perspectives have also been advanced, and our findings support the notion that specific regions have a privileged role in pain representation. Different subregions of mostly ACC, insula, midbrain and cerebellum significantly predict pain on their own. Previously some of these areas have been suggested to be pain specific (a24pr) (Lieberman and Eisenberger 2015) or “fundamental” to pain (OP1) (Segerdahl et al. 2015b) (regions labeled using the scheme of (Glasser et al. 2016)). Substantial evidence indicates cingulate and insular cortex represent parallel pain pathways mediating distinct dimensions of pain experience. Cingulate cortex receives nociceptive afferents from medial dorsal thalamus (Meda et al. 2019; Sikes and Vogt 1992; Yamamura et al. 1996) (predictive here with p = 0.02, uncorrected, fails correction for multiple comparisons), has a history of neurosurgical use in relieving pain affect (Foltz and White 1962; Hurt and Ballantine 1974) and responds preferentially to affective experimental pain manipulations (Rainville et al. 1997; Vogt, Berger, and Derbyshire 2003). Conversely, posterior insula has traditionally been associated with sensory-discriminative dimensions of pain. It receives nociceptive afferents from VPM thalamus (Craig 2014) (significantly pain predictive here), evokes pain upon intracranial stimulation (Mazzola et al. 2012) and insular lesions increase pin-prick pain thresholds (Greenspan, Lee, and Lenz 1999). Convergence between our results and prior studies primarily concerned with regional hypotheses allows our findings to put such studies in a quantified context. We show many brain regions may predict pain with high sensitivity depending on the experimental conditions and sample of subjects observed, but there is substantial unique information elsewhere in the brain (24% more in the full brain vs. best elementary region) which regional interpretations of pain fail to capture.

Resting state network modules also predict pain intensity, especially somatomotor, salience and attention networks. Salience networks respond to painful stimuli (Seeley et al. 2007; Downar et al. 2000; Legrain et al. 2011; Mouraux et al. 2011), leading to interpretations of pain as an orienting response (Baliki and Apkarian 2015). Consistent with this view, we also find default mode network (DMN) deactivation is additionally selected to predict pain across cRSN cross validation folds. The DMN comprises brain areas that are active at rest and implicated in internal monitoring (Andrews-Hanna 2012; Raichle et al. 2001), but proportionally deactivated in response to tasks which engage attention, including pain (Loggia et al. 2012). These results echo network level interpretations of pain, but our brain network parcels of pain are no more pain predictive than elementary region models, correlate with regional pattern maps, and the salience and somatomotor regions selected most frequently show substantial overlap with individually predictive insular and cingulate elementary regions. This indicates that the best a salience network (or any other network) model of pain can hope for is to recapitulate the information already present in the best constituent elementary region.

Multisystem and full brain patterns are the most pain predictive in this study, and provide the greatest model learning capacities. This advantage suggests pain is represented by independent multisystem brain signals, likely constrained to the conjunction of multisystem and full brain models since the full brain shows no unique predictive advantage. Mediation analysis of the full brain predictions indicates mechanisms of temperature and expectation modulation of pain may offer additional guidance. Converging evidence implicates anterolateral, midbrain and cerebellar brain systems in cognitive evaluation, more posterolateral areas in detection of physical stimulus properties, and thalamic and frontal midline regions in both cognitive and physical evaluation of painful stimuli (Kong et al. 2006; Ploner et al. 2011; Ernst et al. 2019; Keltner et al. 2006; Rainville et al. 1997; Seminowicz, Mikulis, and Davis 2004; Ji et al. 2010; Wager et al. 2004; Atlas et al. 2012; Woo et al. 2015), with specific evidence of a double dissociation of expectation and stimulus intensity effects along the anteroposterior axis of the insula (Geuter et al. 2017; Frot, Faillenot, and Mauguière 2014). Although characterizing the diversity of pain information in the brain exceeds the scope of this study, this was precisely the scope of (Atlas et al. 2010) and (Atlas et al. 2014) who use Study 2 and Study 5 (respectively) to identify expectation mediators in dorsolateral PFC, amygdala, pons, and dorsal striatum, thermal stimulus mediators in somatomotor areas and cerebellum, and mediators of both components in dorsal ACC, anterior insula and thalamus. This illustrates the spatial distribution of multidimensional pain signals in our dataset broadly consistent with theoretical conceptions of a pain “neuromatrix”, and lends support to the notion that multisystem models subsume, integrate and expand upon a diverse constellation of unique regional signals.

Several studies suggest traditional methods of functional localization may miss subtle representations within and across brain areas, and that pain representations might be embedded in areas typically associated with unrelated processes (Haxby et al. 2001; Liang et al. 2013). We hoped the full brain hypothesis space would allow our models to capture traditionally overlooked signals, but the failure of full brain models to outperform multisystem models suggests any such exotic signals are redundant with what’s already captured by the multisystem representations. However, capturing full brain representations necessarily requires the greatest model complexity, and therefore is most sensitive to sample size. Additionally, our approach was not optimized to recover fine grained signals which are often obscured by functional heterogeneity across individuals (Haxby et al. 2011). Thus, we cannot rule out the ability of different analytic approaches or larger samples to better leverage full brain information. Whether evoked pain is represented throughout the brain rather than across a multisystem subset of areas remains an open question.

We use MVPA models of pain representation at six spatial scales, estimate their performance, and validate these estimates to show multisystem models best represent pain. We then generalize our conclusions to novel models trained in novel studies, and validate these conclusions as well. In the face of current gyrations between local and global perspectives of pain, our results illustrate the extent to which the neural basis of pain can be localized, that it is best captured by distributed signals spanning multiple brain systems, and provide a blueprint for quantifying information content in the brain across diverse neural phenomena.

## AUTHOR CONTRIBUTIONS

BP, PK and TW conceived of this study and its initial design, however each constituent Study was designed, collected and had GLMs run by different individuals. LA for Studies 2 and 5, SG for Study V3, MJ for Study 3, LK for Study 7, AK for Study 1, ML for Study 6, MR for Study V2, CW for Studies 4 and V1. Additionally TW was involved in the design of all studies except V3. BP performed all statistical and multivariate analyses, and prepared all figures. BP, PK, and TW wrote the report and all authors edited the report and approved the report before submission.

## Notes

https://github.com/canlab/canlab_single_trials

### Competing Interest Statement

The authors have declared no competing interest.

### Summary of Updates

The revision adds links to public repositories where readers can access datasets and code used in the analyses covered by this manuscript. Additionally, some mistakenly omitted acknowledgements were added recognizing support from additional parties.

